# The proton pump inhibitor omeprazole does not promote *Clostridioides difficile* colonization in a murine model

**DOI:** 10.1101/775411

**Authors:** Sarah Tomkovich, Nicholas A. Lesniak, Yuan Li, Lucas Bishop, Madison J. Fitzgerald, Patrick D. Schloss

## Abstract

Proton pump inhibitor (PPI) use has been associated with microbiota alterations and susceptibility to *Clostridioides difficile* infections (CDIs) in humans. We assessed how PPI treatment alters the fecal microbiota and whether treatment promotes CDIs in a mouse model. Mice receiving a PPI treatment were gavaged with 40 mg/kg of omeprazole during a 7-day pretreatment phase, the day of *C. difficile* challenge, and the following 9 days. We found that mice treated with omeprazole were not colonized by *C. difficile*. When omeprazole treatment was combined with a single clindamycin treatment, one cage of mice remained resistant to *C. difficile* colonization, while the other cage was colonized. Treating mice with only clindamycin followed by challenge resulted in *C. difficile* colonization. 16S rRNA gene sequencing analysis revealed that omeprazole had minimal impact on the structure of the murine microbiota throughout the 16 days of omeprazole exposure. These results suggest omeprazole treatment alone is not sufficient to disrupt microbiota resistance to *C. difficile* infection in mice that are normally resistant in the absence of antibiotic treatment.

## Importance

Antibiotics are the primary risk factor for *Clostridioides difficile* infections (CDIs), but other factors may also increase a person’s risk. In epidemiological studies, proton pump inhibitor (PPI) use has been associated with CDI incidence and recurrence. PPIs have also been associated with alterations in the human intestinal microbiota in observational and interventional studies. We evaluated the effects of the PPI omeprazole on the structure of the murine intestinal microbiota and its ability to disrupt colonization resistance to *C. difficile*. We found omeprazole treatment had minimal impact on the murine fecal microbiota and did not promote *C. difficile* colonization. Further studies are needed to determine whether other factors contribute to the association between PPIs and CDIs seen in humans or whether aspects of murine physiology may limit its utility to test these types of hypotheses.

Antibiotics have a large impact on the intestinal microbiome and are a primary risk factor for developing *Clostridioides difficile* infections (CDIs) (1). It is less clear whether other human medications that impact the microbiota also influence *C. difficile* colonization resistance. Multiple epidemiological studies have suggested an association between proton pump inhibitor (PPI) use and incidence or recurrence of CDIs (2–5). There have also been a number of large cohort studies and interventional clinical trials that demonstrated specific alterations in the intestinal microbiome were associated with PPI use (4, 6). PPI-associated microbiota changes have been attributed to the ability of PPIs to increase stomach acid pH which may promote the survival of oral and pathogenic bacteria (4, 6). In human fecal samples, PPI use results in increases in *Enterococcaceae, Lactobacillaceae, Micrococcaceae, Staphylococcaceae* and *Streptococcaceae* and decreases in *Ruminococcaceae* (6–9). Several of these taxa have also been associated with *C. difficile* colonization in humans (10).

Unfortunately, the studies suggesting a link between PPIs and *C. difficile* were retrospective and did not evaluate changes in the microbiome (2, 3, 5). Thus, it is unclear whether the gastrointestinal microbiome changes associated with PPI use explain the association between PPIs and CDIs. Additionally, epidemiological studies have a limited capacity to address potential confounders and comorbidities in patients that were on PPIs and developed CDIs or recurrent CDIs (2, 5). Here, we evaluated the impact of daily PPI treatment with omeprazole on the murine microbiome and susceptibility to *C. difficile* colonization in relation to clindamycin, an antibiotic that perturbs the microbiome enough to allow *C. difficile* to colonize but is mild enough that *C. difficile* is cleared within 10 days (11).

### Murine fecal microbiomes were minimally affected by omeprazole treatment

To test whether omeprazole treatment alters the microbiome and promotes susceptibility to CDIs, we gavaged mice with 40 mg/kg of omeprazole for 7 days before *C. difficile* challenge (Figure 1A). A principle coordinates analysis (PCoA) of the Bray-Curtis distances over the initial 7 days of treatment revealed the bacterial communities of omeprazole-treated mice remained relatively unchanged (Figure 1B). We observed no significant changes in the relative abundance of those taxa previously shown to respond to PPI treatment throughout the course of the 16-day experiment (Figure 1C-D, S1). We also observed no significant changes in relative abundances at the family and genus level over the course of the experiment for the omeprazole-treated mice (all corrected P-values > 0.36). These results demonstrated that the omeprazole treatment alone had a minimal impact on the murine fecal bacterial community after 7 days of pretreatment.

**Figure 1.**
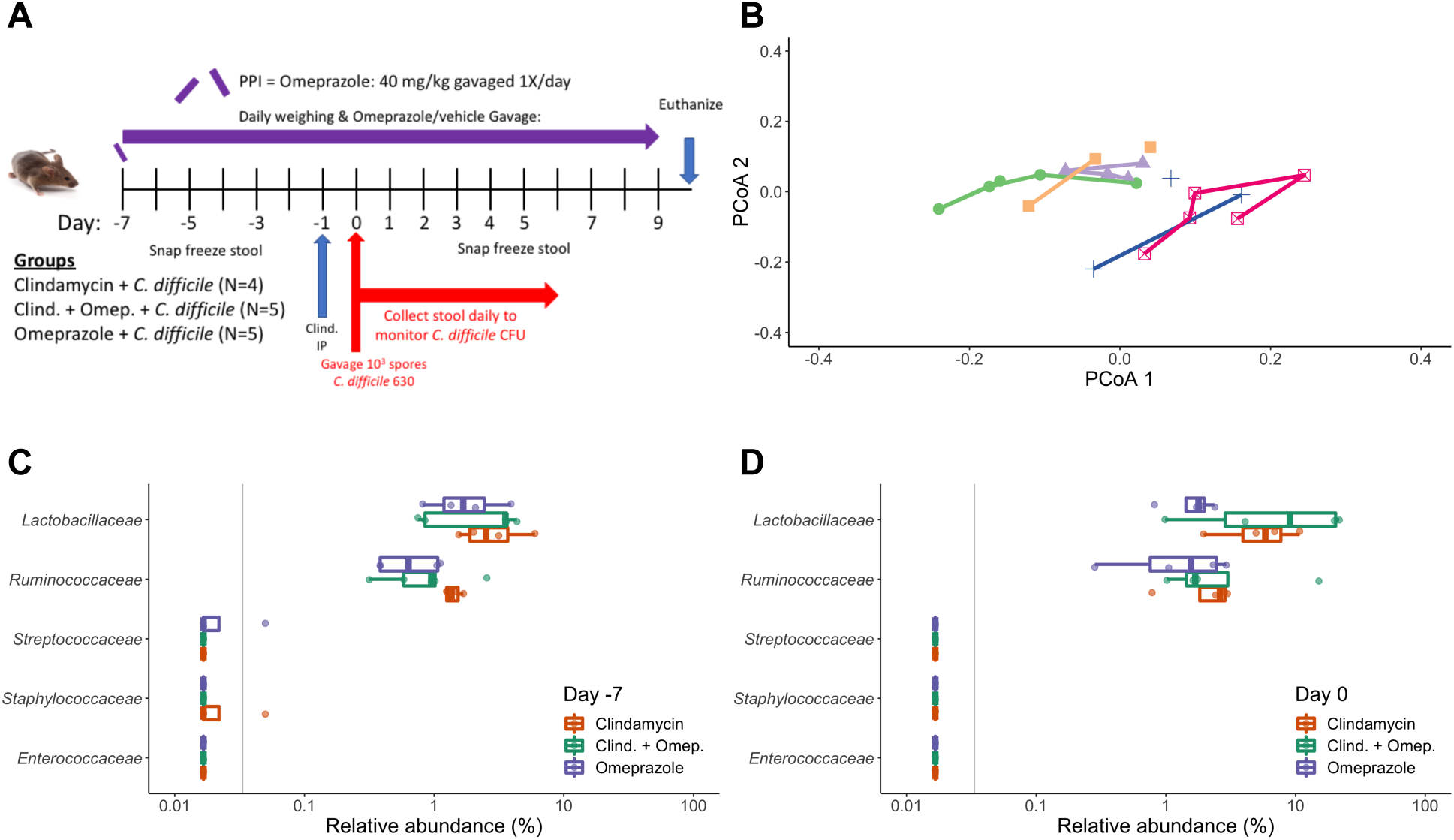
Omeprazole treatment had minimal impact on the murine fecal microbiota. A. Mouse experiment timeline and logistics. The PPI omeprazole was administered throughout the duration of the experiment. Clindamycin was administered 1 day before *C. difficile* challenge on Day 0. Stools for 16S rRNA sequencing analysis were collected on the days that are labeled (Day −7, −5, −3, −1, 0, 1, 2, 3, 4, 5, 7, 9). *C. difficile* CFU in the stool was quantified daily through 6 days post-infection by anaerobic culture. B. Principal Coordinates Analysis (PCoA) of Bray-Curtis distances from stool samples of mice in the omeprazole treatment group during the initial 7 days of the experiment. Each color represents stool samples from the same mouse and lines connect sequentially collected samples. C-D. Relative abundances of families previously associated with PPI use in humans at the start of the experiment (C) and after 7 days of omeprazole treatment (D). Each circle represents an individual mouse. There were no significant differences across treatment groups for any of the identified families in the sequence data at day −7 (all P-values > 0.448) and day 0 (all P-values > 0.137), analyzed by Kruskal-Wallis test with a Benjamini-Hochberg correction for multiple comparisons. For C-D, the grey vertical line indicates the limit of detection.

### Omeprazole treatment did not promote susceptibility to *C. difficile* infection in mice

Next, we examined whether omeprazole treatment altered susceptibility to *C. difficile* infection in mice. After omeprazole treatment or clindamycin treatment, mice were challenged with 10^3^ *C. difficile* 630 spores. Although *C. difficile* colonized the clindamycin-treated mice, it did not colonize the omeprazole-treated mice (Figure 2A). Interestingly, only 1 cage of mice that received both omeprazole and clindamycin were colonized, while the other cage of mice were resistant (Figure 2A). The greatest shifts in bacterial communities occurred in the clindamycin-treated mice (Figure 2B, S2). Regardless of whether the mice became colonized, all of the mice had cleared *C. difficile* within 5 days (Figure 2A), suggesting that omeprazole did not affect the rate of clearance. Our results suggest that omeprazole treatment had no effect on bacterial community resistance to *C. difficile* colonization in mice. Instead most of the differences between the 3 treatment groups appeared to be driven by clindamycin administration (Figure 2C, S2). These findings demonstrated that high dose omeprazole treatment did not promote susceptibility to *C. difficile* colonization.

**Figure 2.**
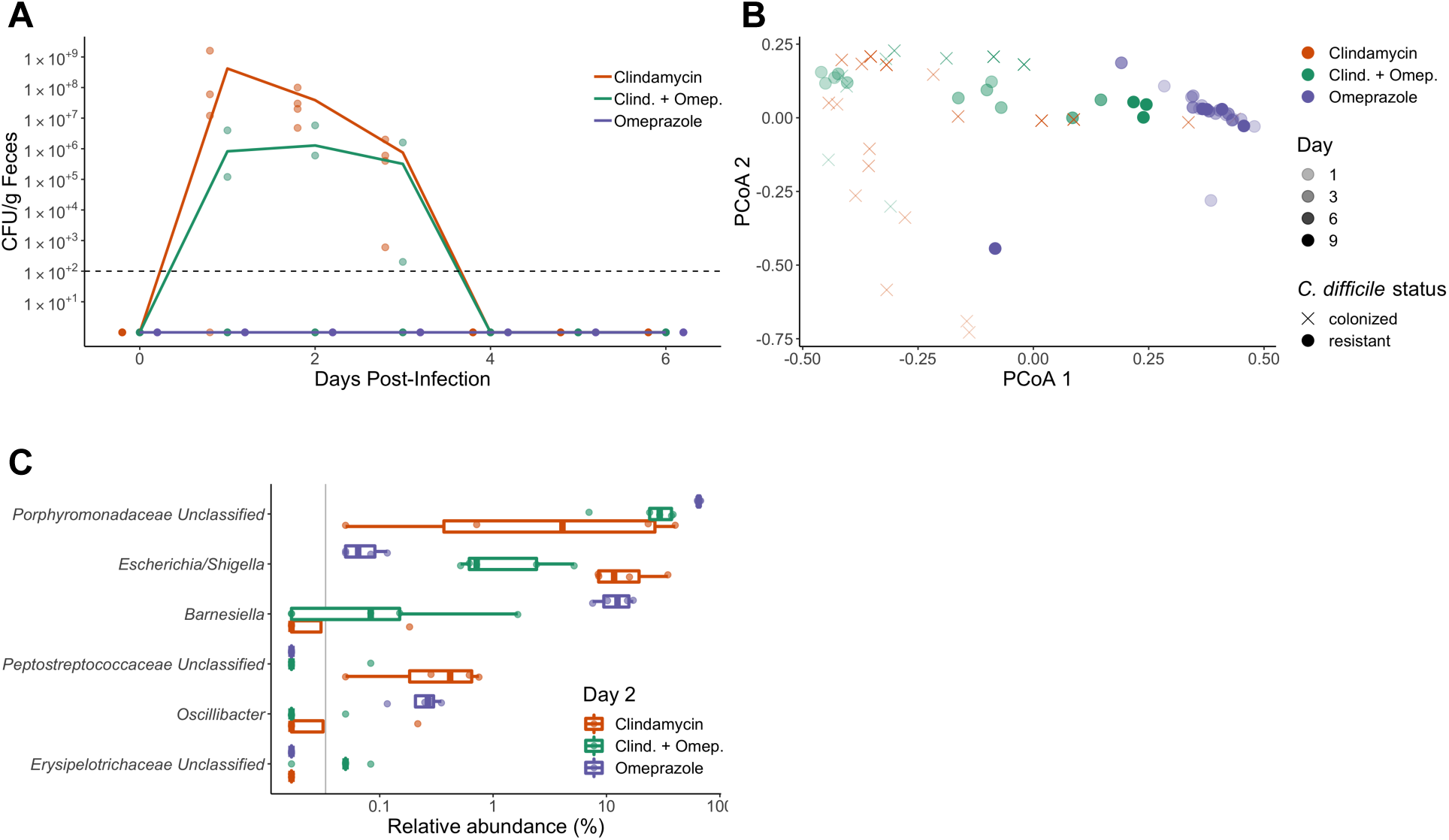
Omeprazole treatment alone does not promote CDIs in mice. A. *C. difficile* CFUs/g stool measured each day post *C. difficile* challenge for clindamycin, clindamycin/omeprazole, and omeprazole-treated mice. Lines represent the mean CFU/g for each treatment group while points represent CFU/g for individual mice within each group. The black dashed line indicates the limit of detection. B. PCoA of of Bray-Curtis distances from stool samples collected after antibiotic treatment (last 9 days of the experiment). Transparency of the symbol corresponds to treatment day. Symbols represent the *C. difficile* colonization status of the mice measured 2 days post-infection. Circles represent resistant mice (*C. difficile* was undetectable in stool samples), while X-shapes represent mice that were colonized with *C. difficile*, although all mice cleared *C. difficile* within 5 days of infection. Omeprazole treated fecal samples primarily cluster together throughout the experiment. C. Genera that vary the most across treatment groups for stool samples collected from mice 2 days post-infection. Data were analyzed by Kruskal-Wallis test, and no P-values were significant after Benjamini-Hochberg correction for multiple comparisons (all P-values > 0.092). The grey vertical line indicates the limit of detection.

### Conclusions

The PPI omeprazole did not meaningfully impact the structure of the gut microbiota and did not promote *C. difficile* infection in mice. Our findings that omeprazole treatment had minimal impact on the fecal microbiome were comparable to another PPI mouse study that indicated the PPI lansoprazole had more of an effect on the small intestinal microbiota compared to the fecal microbiota (12). The same group demonstrated lansoprazole treatment increased the stomach pH in mice (12), which may improve survival of bacteria passing through the stomach. We did not find significant changes in the relative abundances of the taxa observed to be significantly impacted by PPI use in human studies. However, 3 of the human-associated taxa were absent or at low abundance in our mice. Interestingly, other groups examining fecal microbiota communities before and after PPI administration to healthy cats and infants with gastroesophageal reflux disease, found PPIs have minimal effects on fecal bacterial community structures, although there were a few significant changes in specific genera (13, 14). One limitation of our study is that there were only 4-5 mice per group, which may have limited our ability to identify PPI-induced changes in specific bacteria genera. Although our fecal microbiota findings are comparable to what has been shown in another mouse study (12), whether PPI-induced changes in specific bacterial abundances observed in humans play a role in CDIs remains to be determined.

Although several *C. difficile* mouse model studies have shown that PPIs have an effect on CDIs with or without additional antibiotic treatment (15–17), there were insufficient controls to attribute the effect solely to PPI treatment. One group administered 0.5 mg/kg of the PPI lansoprazole daily for 2 weeks to mice and then challenged with *C. difficile* demonstrated that PPI treatment alone resulted in detectable *C. difficile* in the stool 1 week after challenge, however there was detectable *C. difficile* in mice not treated with antibiotics (15, 16). The other mouse study demonstrated antibiotic/esomeprazole-treated mice developed more severe CDIs compared to antibiotic-treated mice, but the researchers did not have a group treated with just esomeprazole for comparison (17). We tested the same high 40 mg/kg PPI dose and expanded pre-treatment to 7 days before challenge to test the impact of omeprazole treatment alone on our CDI mouse model. Additionally, we have previously demonstrated that mice from our breeding colony are resistant to *C. difficile* 630 colonization without antibiotic treatment (18), ensuring there was not already partial susceptibility to *C. difficile* before treatment. The additional controls in our study allowed us to assess the contribution of omeprazole alone to *C. difficile* susceptibility in mice.

Our study also extended previous work examining PPIs and *C. difficile* in mice by incorporating the contribution of the intestinal microbiota. We found omeprazole had no significant impact on bacterial taxa within the murine intestinal microbiota over the 16-day experiment. In contrast to previous work with PPIs (15–17), omeprazole did not alter *C. difficile* colonization resistance in mice. 16S rRNA sequencing suggested that *Streptococcus* and *Enterococcus* are rare genera in our C57BL/6 mouse colony. These two genera could be important contributors to the associations between PPIs and CDIs in humans, and could be a contributing factor to our observation that PPI treatment had no effect on *C. difficile* colonization in our CDI mouse model. While the intestinal microbiomes of both humans and mice are dominated by the *Bacteroidetes* and *Firmicutes* phyla, there are significant differences in the relative abundances of genera that are present and some genera are unique to each mammal (19), differences that may partly explain our results. Gastrointestinal physiological differences, particularly the higher stomach pH in mice (pH 3-4) compared to humans (pH 1) (19) could also explain why omeprazole had a limited impact on the murine microbiome. The microbiota and physiological differences between humans and mice may limit the usefulness of employing mouse models to study the impact of PPIs on the microbiota and CDIs.

Beyond microbiome differences, factors such as age, body mass index, comorbidities, and use of other medications in human studies may also be contributing to the association between PPIs and CDI incidence or recurrence. The type of *C. difficile* strain type could also be an important contributing factor, however our study was limited in that we only tested *C. difficile* 630 (ribotype 012). This study addressed the impact of PPIs with or without antibiotics on a murine model of CDI, and found PPIs did not promote *C. difficile* colonization. The epidemiological evidence linking PPIs to CDIs is primarily from observational studies, which makes determining causality and whether other risk factors play a role challenging (20). Future studies are needed to determine whether age, other comorbidities and bacterial strains that are less common in mice can increase the risk of CDIs or recurrent CDIs when combined with PPI treatment.

## Acknowledgements

This research was supported by NIH grant U01AI12455. We would also like to thank the Unit for Laboratory Animal Medicine at the University of Michigan for maintaining our mouse colony and providing the infrastructure and support for performing our mouse experiments. The authors are also thankful to members of the Schloss lab for helpful discussions throughout the process of designing the experiment, analyzing the results, crafting the figures, and drafting of the manuscript.

## Materials and Methods

### Animals

All mouse experiments were performed with 7- to 12-week-old C57BL/6 male and female mice. Each experimental group of mice was split between 2 cages with 2-3 mice housed per cage and male and female mice housed separately. All animal experiments were approved by the University of Michigan Animal Care and Use Committee (IACUC) under protocol number PRO00006983.

### Drug treatments

Omeprazole (Sigma Aldrich) was prepared in a vehicle solution of 40% polyethylene glycol 400 (Sigma-Aldrich) in phosphate buffered saline. Omeprazole was prepared from 20 mg/mL frozen aliquots and diluted to an 8 mg/mL prior to gavage. All mice received 40 mg/kg omeprazole (a dose previously used in mouse experiments (17)) or vehicle solution once per day through the duration of the experiment with treatment starting 7 days before *C. difficile* challenge (Figure 1A). Although the omeprazole dose administered to mice is higher than the recommended dose for humans, omeprazole has a shorter half-life in mice compared to humans (21) and lacks an enteric coating (22). One day prior to *C. difficile* challenge, 2 groups of mice received an intraperitoneal injection of 10 mg/kg clindamycin or sterile saline vehicle (11). All drugs were filter sterilized through a 0.22 micron syringe filter before administration to animals.

### *C. difficile* infection model

Mice were challenged with *C. difficile* 630 seven days after the start of omeprazole treatment and one day after clindamycin treatment. Mice were challenged with 10^3^ spores in ultrapure distilled water as described previously (11). Stool samples were collected for 16S rRNA sequencing or *C. difficile* CFU quantification throughout the duration of the experiments at the indicated timepoints (Figure 1A). Samples for 16S rRNA sequencing were flash frozen in liquid nitrogen and stored at −80°C until DNA extraction, while samples for CFU quantification were transferred into an anaerobic chamber and serially diluted in PBS. Diluted samples were plated on TCCFA (taurocholate, cycloserine, cefoxitin, fructose agar) plates and incubated at 37°C for 24 hours under anaerobic conditions to quantify *C. difficile* CFU.

### 16S rRNA gene sequencing

DNA for 16S rRNA gene sequencing was extracted from 10-50 mg fecal pellet from each mouse using the DNeasy Powersoil HTP 96 Kit (Qiagen) and an EpMotion 5075 automated pipetting system (Eppendorf). The 16S rRNA sequencing library was prepared as described previously (23). In brief, the ZymoBIOMICS™ Microbial Community DNA Standard (Zymo, CA, USA) was used as a mock community (24) and water was used as a negative control. The V4 hypervariable region of the 16S rRNA gene was amplified with Accuprime Pfx DNA polymerase (Thermo Fisher Scientific) using previously described custom barcoded primers (23). The 16S rRNA amplicon library was sequenced with the MiSeq (Illumina). Amplicons were cleaned up and normalized with the SequalPrep Normalization Plate Kit (ThermoFisher Scientific) and pooled amplicons were quantified with the KAPA library quantification kit (KAPA Biosystems).

### 16S rRNA gene sequence analysis

mothur (v1.40.5) was used for all sequence processing steps (25) using a previously published protocol (23). In brief, forward and reverse reads for each sample were combined and low-quality sequences and chimeras were removed. Duplicate sequences were merged, before taxonomy assignment using a modified version (v16) of the Ribosomal Database Project reference database (v11.5) with an 80% cutoff. Operational taxonomic units (OTUs) were assigned with the opticlust clustering algorithm using a 97% similarity threshold. To adjust for uneven sequencing across samples, all samples were rarefied to 3,000 sequences, 1,000 times. PCoAs were generated based on Bray-Curtis distance. R (v.3.5.1) was used to generate figures and perform statistical analysis.

### Statistical Analysis

To test for differences in relative abundances in families and genera across our 3 different treatment groups at different timepoints (Clindamycin, Clindamycin + Omeprazole, and Omeprazole on Day −7, 0, 2, and 9) or within the Omeprazole treatment group across 3 timepoints (Day −7, 0, and 9), we used a Kruskal-Wallis test with a Benjamini-Hochberg correction for multiple comparisons.

### Code availability

The code for all sequence processing and analysis steps as well as a Rmarkdown version of this manuscript is available at https://github.com/SchlossLab/Tomkovich_PPI_mSphere_2019.

### Data availability

The 16S rRNA sequencing data have been deposited in the NCBI Sequence Read Archive (Accession no. PRJNA554866).

## Figures

**Figure S1.**
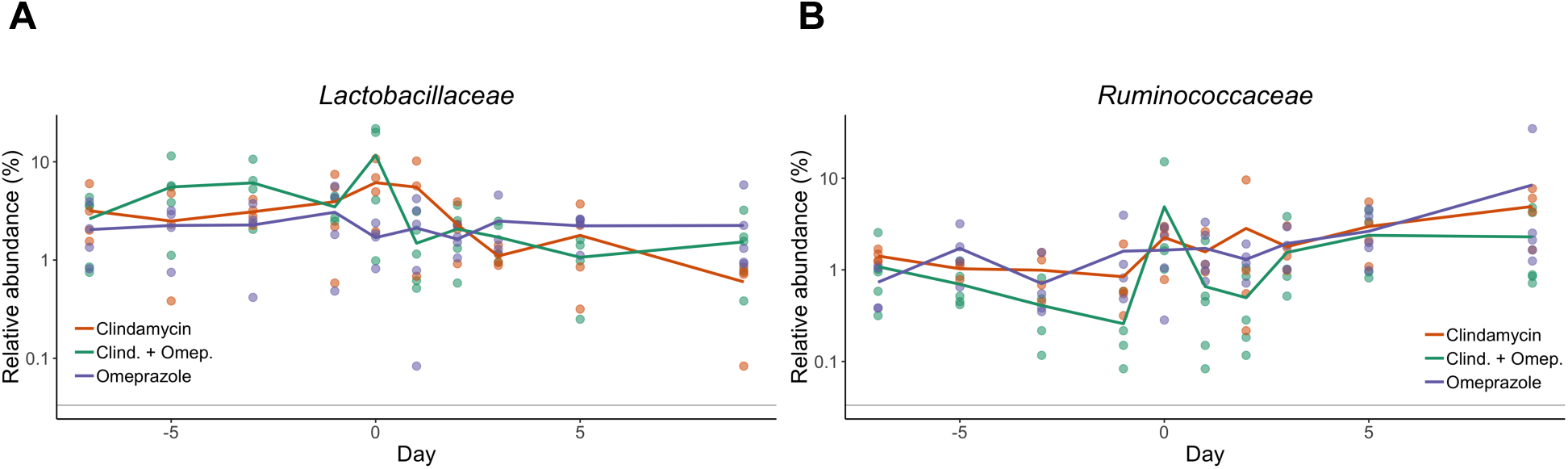
Families within omeprazole treated mice fluctuate over time with no overall trend in either direction. Relative abundance over time for *Lactobacillaceae* (A) and *Ruminococcaceae* (B), 2 of the PPI-associated families from human PPI studies across all 3 treatment groups. Each point represents the relative abundance for an individual mouse stool sample, while the lines represent the mean relative abundances for each treatment group of mice. The grey horizontal lines indicate the limit of detection.

**Figure S2.**
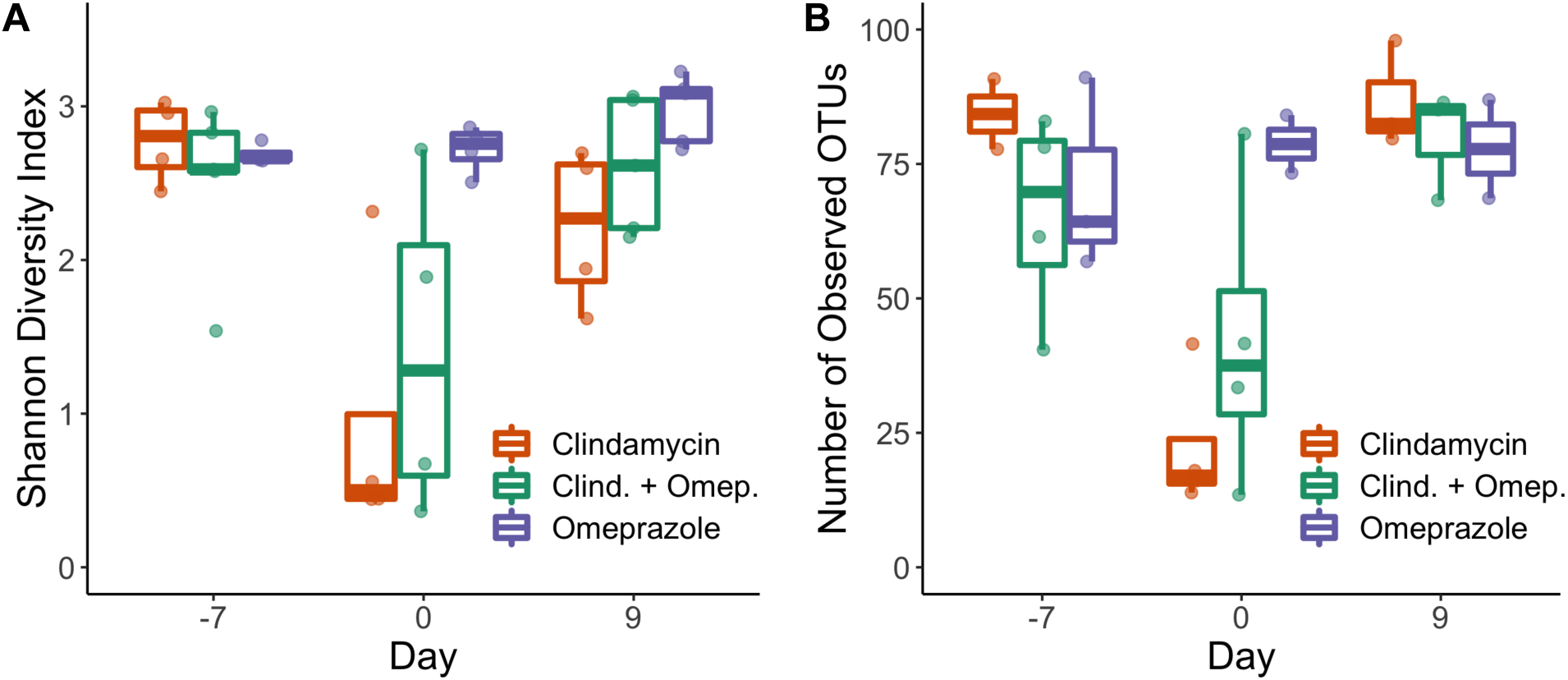
Microbiota diversity and richness decrease with antibiotic treatment but remain relatively constant with omeprazole treatment. Boxplots of the Shannon Diversity Index values (A) and number of observed OTUs (B) for each group of mice over 3 timepoints (Day −7, 0, and 9). Each circle represents the value for a stool sample from an individual mouse.

## References

1. Schubert AM, Sinani H, Schloss PD. 2015. Antibiotic-induced alterations of the murine gut microbiota and subsequent effects on colonization resistance against *Clostridium difficile*. mBio 6.

2. Tariq R, Singh S, Gupta A, Pardi DS, Khanna S. 2017. Association of gastric acid suppression with recurrent *Clostridium difficile* infection: A systematic review and meta-analysis. JAMA internal medicine 177:784–791.

3. Nehra AK, Alexander JA, Loftus CG, Nehra V. 2018. Proton pump inhibitors: Review of emerging concerns, pp. 240–246. In Mayo clinic proceedings. Elsevier.

4. Naito Y, Kashiwagi K, Takagi T, Andoh A, Inoue R. 2018. Intestinal dysbiosis secondary to proton-pump inhibitor use. Digestion 97:195–204.

5. Elias E, Targownik LE. 2019. The clinician’s guide to proton pump inhibitor related adverse events. Drugs 79:715–731.

6. Imhann F, Vila AV, Bonder MJ, Manosalva AGL, Koonen DP, Fu J, Wijmenga C, Zhernakova A, Weersma RK. 2017. The influence of proton pump inhibitors and other commonly used medication on the gut microbiota. Gut Microbes 8:351–358.

7. Imhann F, Bonder MJ, Vila AV, Fu J, Mujagic Z, Vork L, Tigchelaar EF, Jankipersadsing SA, Cenit MC, Harmsen HJM, Dijkstra G, Franke L, Xavier RJ, Jonkers D, Wijmenga C, Weersma RK, Zhernakova A. 2015. Proton pump inhibitors affect the gut microbiome. Gut 65:740–748.

8. Freedberg DE, Toussaint NC, Chen SP, Ratner AJ, Whittier S, Wang TC, Wang HH, Abrams JA. 2015. Proton pump inhibitors alter specific taxa in the human gastrointestinal microbiome: A crossover trial. Gastroenterology 149:883–885.e9.

9. Maier L, Pruteanu M, Kuhn M, Zeller G, Telzerow A, Anderson EE, Brochado AR, Fernandez KC, Dose H, Mori H, others. 2018. Extensive impact of non-antibiotic drugs on human gut bacteria. Nature 555:623.

10. Schubert AM, Rogers MAM, Ring C, Mogle J, Petrosino JP, Young VB, Aronoff DM, Schloss PD. 2014. Microbiome data distinguish patients with *Clostridium difficile* infection and non-*C. difficile*-associated diarrhea from healthy controls. mBio 5.

11. Jenior ML, Leslie JL, Young VB, Schloss PD. 2018. *Clostridium difficile* alters the structure and metabolism of distinct cecal microbiomes during initial infection to promote sustained colonization. mSphere 3.

12. Yasutomi E, Hoshi N, Adachi S, Otsuka T, Kong L, Ku Y, Yamairi H, Inoue J, Ishida T, Watanabe D, Ooi M, Yoshida M, Tsukimi T, Fukuda S, Azuma T. 2018. Proton pump inhibitors increase the susceptibility of mice to oral infection with enteropathogenic bacteria. Digestive Diseases and Sciences 63:881–889.

13. Schmid SM, Suchodolski JS, Price JM, Tolbert MK. 2018. Omeprazole minimally alters the fecal microbial community in six cats: A pilot study. Frontiers in Veterinary Science 5.

14. Castellani C, Singer G, Kashofer K, Huber-Zeyringer A, Flucher C, Kaiser M, Till H. 2017. The influence of proton pump inhibitors on the fecal microbiome of infants with gastroesophageal refluxA prospective longitudinal interventional study. Frontiers in Cellular and Infection Microbiology 7.

15. Kaur S, Vaishnavi C, Prasad KK, Ray P, Kochhar R. 2007. Comparative role of antibiotic and proton pump inhibitor in experimental *Clostridium difficile* infection in mice. Microbiology and Immunology 51:1209–1214.

16. Kaur S, Vaishnavi C, Prasad KK, Ray P, Kochhar R. 2011. Effect of lactobacillus acidophilus & epidermal growth factor on experimentally induced *Clostridium difficile* infection. The Indian journal of medical research 133:434.

17. Hung Y-P, Ko W-C, Chou P-H, Chen Y-H, Lin H-J, Liu Y-H, Tsai H-W, Lee J-C, Tsai P-J. 2015. Proton-pump inhibitor exposure aggravates *Clostridium difficile*–associated colitis: Evidence from a mouse model. The Journal of infectious diseases 212:654–663.

18. Jenior ML, Leslie JL, Young VB, Schloss PD. 2017. *Clostridium difficile* colonizes alternative nutrient niches during infection across distinct murine gut microbiomes. mSystems 2.

19. Hugenholtz F, Vos WM de. 2018. Mouse models for human intestinal microbiota research: A critical evaluation. Cellular and Molecular Life Sciences 75:149–160.

20. Eze P, Balsells E, Kyaw MH, Nair H. 2017. Risk factors for *Clostridium difficile* infections – an overview of the evidence base and challenges in data synthesis. Journal of Global Health 7.

21. Regårdh C-G, Gabrielsson M, Hoffman K-J, Löfberg I, Skånberg I. 1985. Pharmacokinetics and metabolism of omeprazole in animals and man - an overview. Scandinavian Journal of Gastroenterology 20:79–94.

22. Llorente C, Jepsen P, Inamine T, Wang L, Bluemel S, Wang HJ, Loomba R, Bajaj JS, Schubert ML, Sikaroodi M, Gillevet PM, Xu J, Kisseleva T, Ho SB, DePew J, Du X, Sørensen HT, Vilstrup H, Nelson KE, Brenner DA, Fouts DE, Schnabl B. 2017. Gastric acid suppression promotes alcoholic liver disease by inducing overgrowth of intestinal enterococcus. Nature Communications 8.

23. Kozich JJ, Westcott SL, Baxter NT, Highlander SK, Schloss PD. 2013. Development of a dual-index sequencing strategy and curation pipeline for analyzing amplicon sequence data on the MiSeq illumina sequencing platform. Applied and Environmental Microbiology 79:5112–5120.

24. Sze MA, Schloss PD. 2019. The impact of DNA polymerase and number of rounds of amplification in PCR on 16S rRNA gene sequence data. mSphere 4.

25. Schloss PD, Westcott SL, Ryabin T, Hall JR, Hartmann M, Hollister EB, Lesniewski RA, Oakley BB, Parks DH, Robinson CJ, Sahl JW, Stres B, Thallinger GG, Horn DJV, Weber CF. 2009. Introducing mothur: Open-source, platform-independent, community-supported software for describing and comparing microbial communities. Applied and Environmental Microbiology 75:7537–7541.

